# Sensory neuropathy-causing mutations in ATL3 cause aberrant ER membrane tethering

**DOI:** 10.1101/192484

**Authors:** Michiel Krols, Sammy Detry, Bob Asselbergh, Leonardo Almeida-Souza, Anna Kremer, Saskia Lippens, Riet de Rycke, Vicky De Winter, Franz-Josef Müller, Ingo Kurth, Harvey T. McMahon, Savvas N. Savvides, Vincent Timmerman, Sophie Janssens

**Author notes:** **Address of correspondence:** Prof. Dr. Sophie Janssens, PhD, Laboratory of ER stress and Inflammation, VIB-UGhent, Center for Inflammation Research, Technologiepark 927, B-9052 Zwijnaarde, Belgium, Prof. Dr. Vincent Timmerman, PhD, Peripheral Neuropathy Research Group, University of Antwerp - CDE, Parking P4, Building V, Room 1.30, Universiteitsplein 1, 2610 Antwerpen, Belgium. These authors contributed equally (Michiel Krols, Sammy Detry, Bob Asselbergh, Vincent Timmerman, Sophie Janssens).

## Abstract

The ER is a complex network of sheets and tubules that is continuously being remodeled. The relevance of this membrane dynamics is underscored by the fact that mutations in Atlastins (ATL), the ER fusion proteins in mammals, cause neurodegeneration. How defects in this process disrupt neuronal homeostasis is largely unknown. Here we show by EM volume reconstruction of transfected cells, neurons and patient fibroblasts that the HSAN-causing ATL3 mutants promote aberrant ER tethering hallmarked by bundles of laterally attached ER tubules. *In vitro*, these mutants cause excessive liposome tethering, recapitulating the results in cells. Moreover, ATL3 variants retain their dimerization-dependent GTPase activity, but are unable to promote membrane fusion, suggesting a defect on an intermediate step of the ATL3 functional cycle. Our data therefore show that the effects of ATL3 mutations on ER network organization stretch beyond a loss of fusion, shedding a new light on neuropathies caused by atlastin defects.

## INTRODUCTION

The endoplasmic reticulum (ER) is an intricate network of sheets and tubules that spreads throughout the cytoplasm and is continuous with the nuclear envelope. The distinct morphologies of the ER subdomains are determined by a tug of war between proteins promoting the high membrane curvature found in ER tubules and sheet edges on the one hand, and proteins segregating to and stabilizing ER sheets on the other. Adding to the complexity of this organelle, the network is continuously being remodeled through the extension and retraction of tubules, tubule fusions and ring closures (Westrate et al., 2015).

In metazoans, homotypic fusion of ER tubules is mediated by a class of membrane-bound dynamin-like GTPases known as Atlastins (ATLs) (Hu et al., 2009; Orso et al., 2009). The ATL proteins are comprised of a large N-terminal cytoplasmic domain, two closely spaced transmembrane helices that anchor the protein in the ER membrane, and a C-terminal amphipathic helix. The N-terminal domain consists of a GTPase domain (G domain) connected via a flexible linker to a three-helix bundle (3HB) middle domain (Bian et al., 2011; Byrnes and Sondermann, 2011; Liu et al., 2012). Homotypic ER membrane fusion is mediated by the GTP-dependent homodimerization of ATLs on opposing membranes, promoting tight membrane tethering and eventually membrane fusion (Byrnes and Sondermann, 2011; Byrnes et al., 2013; Hu and Rapoport, 2016; Liu et al., 2015; O’Donnell et al., 2017; Saini et al., 2014)(see Fig. S1). Although the actual fusion mechanism has not been entirely elucidated, it is clear that the ATL transmembrane domains and the C-terminal tail are crucial for membrane fusion, presumably by destabilizing the membranes to promote lipid mixing (Liu et al., 2012; Moss et al., 2011).

Mammals have three ATLs, which share high sequence homology but are differentially expressed. ATL1 is predominantly expressed in the brain, whereas ATL2 and ATL3 show a more ubiquitous expression pattern (Rismanchi et al., 2008). In addition, the three ATL proteins appear to have different fusogenic capacities, with ATL1 being the stronger fusogen and ATL3 a much weaker one (Hu et al., 2015; O’Donnell et al., 2017). In line with this, selective loss of ATL3 has very little effect on ER morphogenesis, whereas ATL1 alone is sufficient to rescue the loss of both ATL2 and ATL3 in COS-7 cells (Hu et al., 2015).

Mutations in ATL1 have been identified in patients suffering from degeneration of upper motor neurons (hereditary spastic paraplegia, HSP) as well as peripheral sensory neurons (hereditary sensory and autonomic neuropathy, HSAN) (Guelly et al., 2011; Zhao et al., 2001). More recently two autosomal dominant missense-mutations in ATL3 were found to cause sensory neurodegeneration: p.Tyr192Cys and p.Pro338Arg, hereafter referred to as ATL3^Y192C^ and ATL3^P338R^ (Fischer et al., 2014; Kornak et al., 2014). HSP patients show progressive spasticity in the lower limbs whereas individuals affected by HSAN have a characteristic loss of pain perception and variable degrees of autonomic dysfunction. Common to both diseases is a degeneration of neurons with particularly long axons. Interestingly, mutations in additional ER shaping proteins such as REEP1, REEP2, Spastin, and Reticulon 2 cause similar phenotypes (Hübner and Kurth, 2014), clearly showing that maintaining the dynamic structure of the ER throughout these very long axons is pivotal for neuronal survival. What is less clear, however, is how defects in ER shaping underlie axonal degeneration. We therefore investigated whether disease-causing mutations in ATL3 alter its functionality and how this affects ER shaping and dynamics. Here, we reveal for the very first time evidence for aberrant ER membrane tethering caused by ATL3 mutations. Using volume electron microscopy, we provide unprecedented insights in the structure of the ER tangles resulting from this aberrant tethering in cells expressing mutant ATL3. Given the redundancy of the three atlastin proteins, this aberrant ER membrane tethering likely represents a major contribution to the axonal degeneration observed in patients.

## RESULTS

### Mutations in ATL3 disrupt ER morphogenesis

We recently uncovered two novel mutations in ATL3 to be causative for HSAN and with this finding added ATL3 to a growing list of ER shaping proteins involved in neurodegeneration (Fischer et al., 2014; Kornak et al., 2014). To study the impact of the ATL3 Y192C and P338R mutations on ER morphology, we imaged COS-7 cells co-expressing mRuby-KDEL as a luminal ER marker (Kredel et al., 2009) and GFP-tagged wild-type (WT) and mutant versions of ATL3 (Fig. S2). ATL3^WT^ was targeted to the ER where it was present mostly in the highly-curved membranes of ER tubules (Fig. 1A). ATL3^Y192C^ or ATL3^P338R^ similarly localized to the tubular ER (Fig. 1A). However, while ATL3^WT^ spread evenly throughout the ER network, mutant ATL3 appeared in patches along the tubules in the peripheral ER and was present mostly in an ER region near the nucleus (Fig. 1A). Similar phenotypes were observed in HeLa cells (data not shown). The peripheral ER network morphology was dramatically altered in cells expressing mutant ATL3. While the ER in control cells formed a highly interconnected tubular network in the periphery of the cell (Fig. 1B), cells transfected with ATL3^Y192C^ or ATL3^P338R^ displayed a peripheral ER network that was far more disperse (Fig. 1B), as quantified by the significant decrease in density of threeway junctions per cell area (Fig. 1B,C). Concordantly, expression of either mutant ATL3 resulted in larger zones within the cell periphery that were devoid of ER tubules, defined as ER sparsity (Fig. 1B,D).

**Fig. 1:**
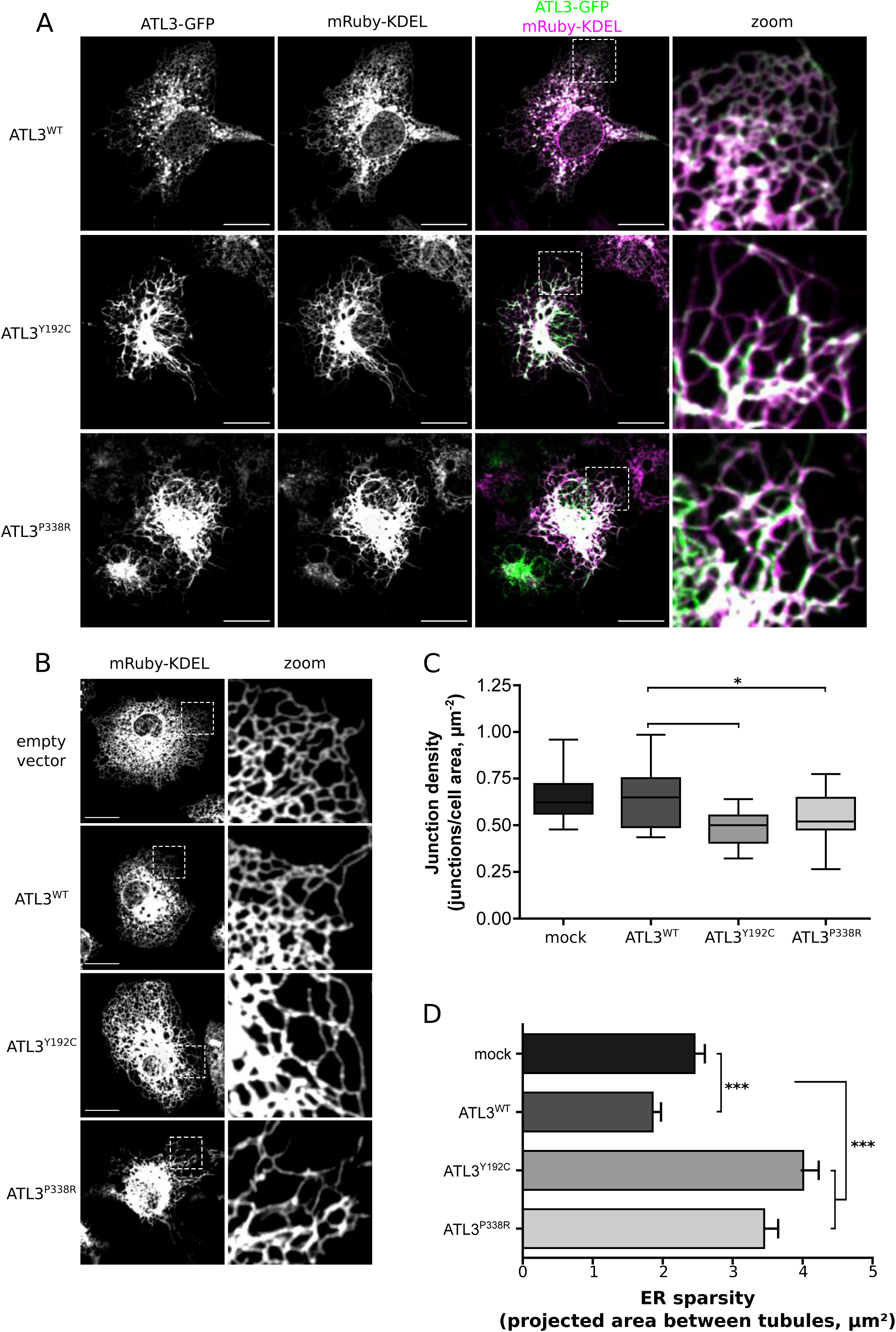
Mutations in ATL3 abate ER network connectivity. A) COS-7 cells were transfected with ATL3-GFP (green) and mRuby-KDEL as an ER marker (magenta). A higher magnification image of the boxed area is shown in the right panel. Scale bar = 10µm. B) The mRubyKDEL ER-marker was imaged in COS-7 cells transfected with empty vector, ATL3^WT^, ATL3^Y192C^ or ATL3^P338R^, as indicated. A higher magnification of the boxed area is shown. Scale bar = 10µm. C) The number of threeway junctions per cell area (junction density) was quantified as a measure of peripheral ER connectivity. Boxplots show the median and the 1^st^ and 3^rd^ quartiles. Whiskers indicate the minimum/maximum values. D) ER sparsity was measured as the average surface area of the polygons formed by ER tubules. Mean +/-SEM is plotted. Statistical analysis in C) and D) was performed using one-way Anova followed by Tukey’s multiple comparisons test. *: P<0.05; ***: P<0.001; n = 22-32 cells per genotype. See also Fig. S2-3.

Besides ER morphogenesis, several reports have suggested a role for ATLs in the Golgi apparatus (Namekawa et al., 2007; Rismanchi et al., 2008). Neither WT nor mutant ATL3 were found to localize to the Golgi complex of HeLa or COS-7 cells, and overexpression of mutant ATL3 was not found to alter Golgi apparatus morphology in these cells (Fig. S3), indicating that a disruption of the ER tubular network is the main hallmark of these mutations in ATL3.

### ER continuity is maintained in cells overexpressing mutant ATL3

A key feature of the ER is that despite the continuous membrane rearrangements, it maintains a single continuous lumen. In a recently reported *in vitro* reconstitution model of the ER, blocking ATL-mediated fusion resulted in fast fragmentation of the network (Powers et al., 2017). Given the severe effect of the mutations in ATL3 on ER morphology, we examined whether ER continuity was maintained in cells expressing mutant ATL3. Therefore, we co-expressed a photoactivatable GFP targeted to the ER lumen (PA-GFP-KDEL) (Jones et al., 2009) in cells expressing mCherry-tagged ATL3. Photo-induced activation of this fluorescent protein allows tracking PA-GFP-KDEL as it pervades from the site of activation throughout the ER network, and thus allows assessing its luminal continuity (Video 1). In addition, the rate at which the fluorophores spreads throughout the ER can be used as a measure of network connectivity (Fig. S4). In cells expressing ATL3^WT^, the activated PA-GFP rapidly diffuses from its site of activation throughout the ER network (Fig. 2A). Photo-activation in similar regions of the ER in mutant ATL3 expressing cells comparably resulted in the spread of the activated fluorophores throughout the entire network, indicating that luminal continuity is not disrupted in these cells (Fig. 2A). However, in agreement with a loss of threeway junctions, analysis of the rate at which the fluorescence intensity increased at discrete distances from the site of activation shows a significantly slower emergence of activated PA-GFP in distal regions of cells expressing ATL3^Y192C^ or ATL3^P338R^ compared to cells expressing ATL3^WT^ (Fig. 2B and Fig. S4). While other characteristics of network topology may also contribute to an altered diffusion speed, this slowed diffusion through the ER network is in agreement with a failure to establish an intricately connected ER network in cells overexpressing mutant ATL3 (Fig. 1). Combined, these findings show that mutations in ATL3 affect the formation of threeway junctions without causing network fragmentation.

**Fig. 2:**
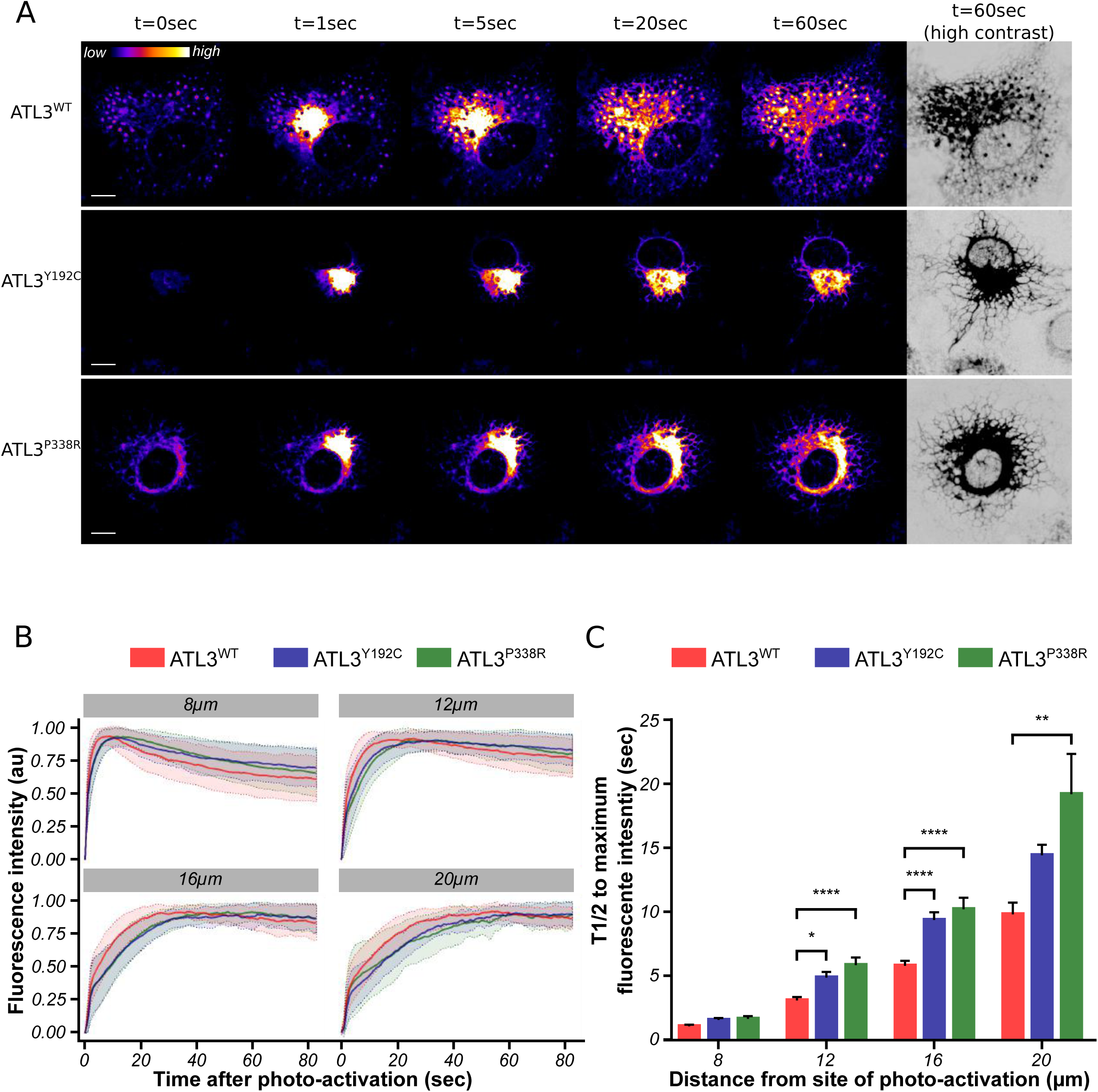
ER continuity is not lost in ATL3 mutant cells. ER-targeted photo-activatable GFP (PA-GFP-KDEL) was imaged at 1 frame per second in COS-7 cells transfected with ATL3^WT^, ATL3^Y192C^ or ATL3^P338R^, as indicated (see Video 1). A) PA-GFP-KDEL was activated in a small area next to the nucleus, after which the activated fluorophores diffuse through the lumen of the entire ER network. Fluorescence intensity color code of the PA-GFP-KDEL is shown on the top left. The frame just prior to activation is indicated as t=0sec and the frame immediately following activation as t=1sec. To clearly show the peripheral ER network, the last frame is also shown as a high-contrast, inverted grayscale image. Scale bar = 10µm. B) The increase in fluorescence intensity over time was measured in regions of interest at fixed distances (8, 12, 16 and 20μm) from the site of activation (see also Fig. S4). Ribbon plots show the mean fluorescence intensity ± SEM for cells expressing ATL3^WT^ (red), ATL3^Y192C^ (blue) and ATL3^P338R^ (green). C) The half-time to reach maximum intensity was extracted by fitting an exponential curve to the phase of fluorescence increase in each region of interest, as highlighted in Fig. S4. Mean ± SEM is plotted and statistical analysis was performed using two-way Anova followed by Tukey’s multiple comparisons test. *: P<0.05; **: P<0.01; ****: P<0.0001; n = 40-75 regions of interest per distance in 10-17 cells per genotype.

### Overexpression of mutant ATL3 causes a defect in ER membrane fusion

Threeway junctions represent the nodes that interconnect the tubular ER and are established through homotypic ER membrane fusion. The loss of junctions in ATL3 mutant cells suggests that mutations in ATL3 disrupt ER membrane fusion. Therefore, live cell imaging of COS-7 cells expressing ATL3-GFP and mRuby-KDEL as an ER marker was performed to monitor events in which a newly forming tubule extends from an existing one and makes contact with an opposing tubule (Fig. 3A). When this contact resulted in a novel junction, it was scored as a successful ER membrane fusion event (Fig. 3A, Video 2). In contrast, when this interaction failed to establish a threeway junction and resulted in retraction of the extending tubule, it was considered a failed fusion event (Fig. 3A, Video 3). Compared to control cells, overexpression of ATL3^WT^ significantly increased the ER membrane fusion success rate from 67 ± 2% to 78 ± 2% (Fig. 3B). Expression of mutant ATL3 on the other hand resulted in a dramatic reduction of this success rate to 23 ± 2% in the case of ATL3^Y192C^ and 40 ± 2% in the case of ATL3^P338R^ (Fig. 3B). Hence, the efficiency of ER membrane fusion is impaired dramatically when ATL3 carrying the HSAN-causing mutations is expressed, indicating that the mutations cause a defect in the mechanism whereby ATL3 induces membrane fusion.

**Fig. 3:**
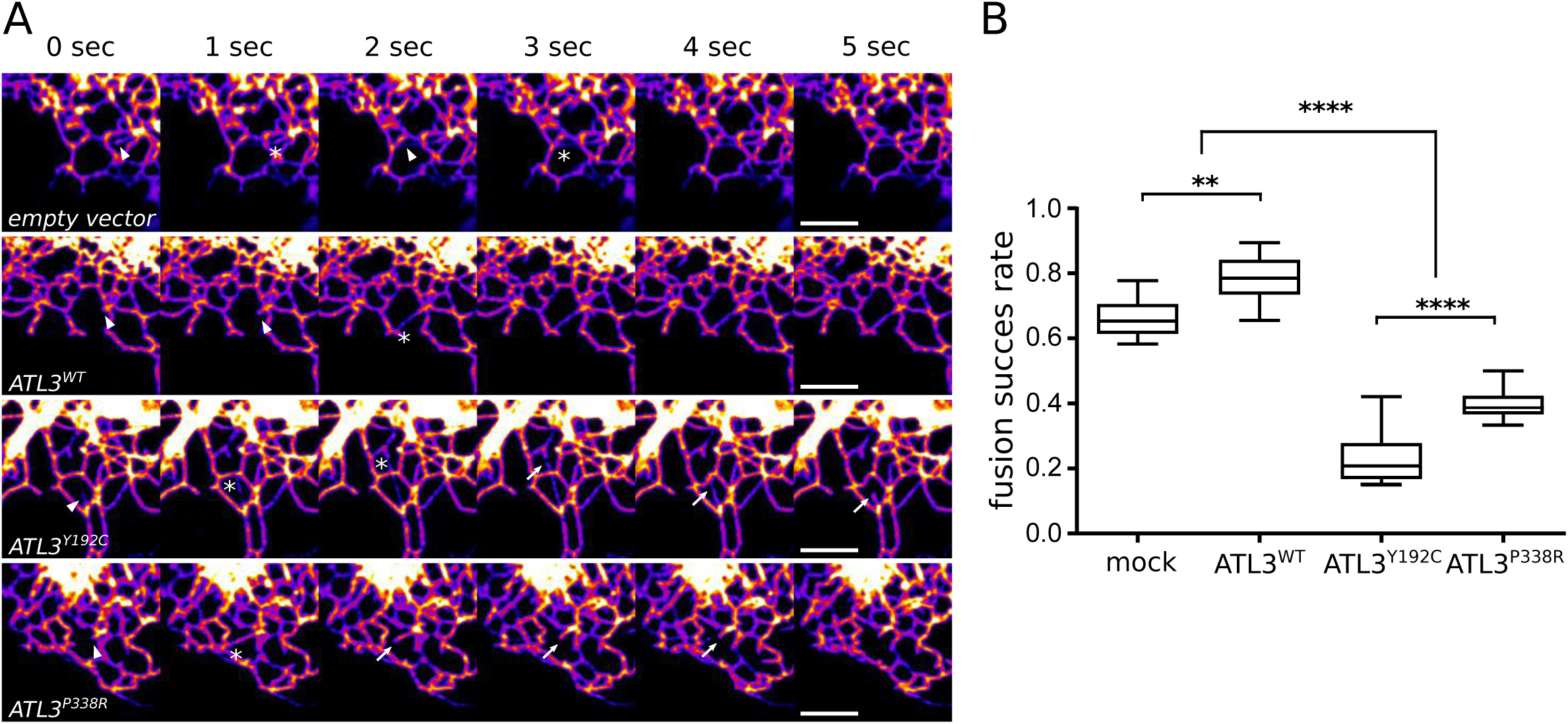
ATL3 mutations cause loss of ER fusion efficiency. Live cell imaging was performed to examine the effect of mutant ATL3 expression on ER tubule fusion. A) The mRubyKDEL ER-marker was imaged at a frame rate of 1 sec^−1^ in COS-7 cells transfected with empty vector, ATL3^WT^, ATL3^Y192C^ or ATL3^P338R^, as indicated (see Videos 2,3). Arrowheads mark the tips of extending tubules. An encounter between the extending tubule and another one is marked by an asterisk. Arrows mark the tip of a retracting tubule. Scale bar = 10µm. B) The success rate of tubule fusion was determined as the fraction of tubule encounter events (as marker by asterisks in A) that result in the formation of stable junctions. Events where tubules retract after an encounter (as indicated by arrows in A) were counted as failed fusion events. Boxplots and whiskers indicate mean and minimum/maximum values. Statistical analysis was performed using one-way Anova, followed by Tukey’s multiple comparisons test. **: P<0.01; ****: P<0.0001; n = 8-11 cells per genotype.

### Mutations in ATL3 interfere with GTP hydrolysis-dependent dimerization

To probe the possible mechanistic consequences of the Y192C and P338R mutations in the functional cycle of ATL3, we compared the biochemical and biophysical properties of the soluble domain of ATL3^WT^, ATL3^Y192C^ and ATL3^P338R^ encompassing the GTPase domain, the linker and the 3HB middle domain (Fig. S5A). Prior structural studies on ATL1 had revealed the existence of at least two distinct conformations of ATL1, termed “extended” and “crossover” (Fig. S1), which were hypothesized to correspond to the prefusion and postfusion states of ATL1, respectively (Bian et al., 2011; Byrnes and Sondermann, 2011). In the extended conformation, for which a crystal structure of ATL3 was also recently solved (O’Donnell et al., 2017), the alpha-helical 3HB middle domain packs tightly against the G domain, enforced mainly by hydrophobic interactions (Fig. 4A, S1). Interestingly, the Y192 and P338 residues in ATL3, affected in HSAN patients, localize to this hydrophobic core (Fig. 4A). At the same time, these amino acid residues pack against the hinge region that relays the drastic structural transitioning of ATL3 into its crossover conformation upon hydrolysis of GTP (Fig. 4A). It is thus clear that both mutations affect residues that are positioned within a functional switch region in ATL3 (Fig. 4A). The functional relevance of these residues is further underscored by the fact that mutations at the equivalent positions in the homologous ATL1 cause HSP (de Bot et al., 2013; McCorquodale et al., 2011).

**Fig. 4:**
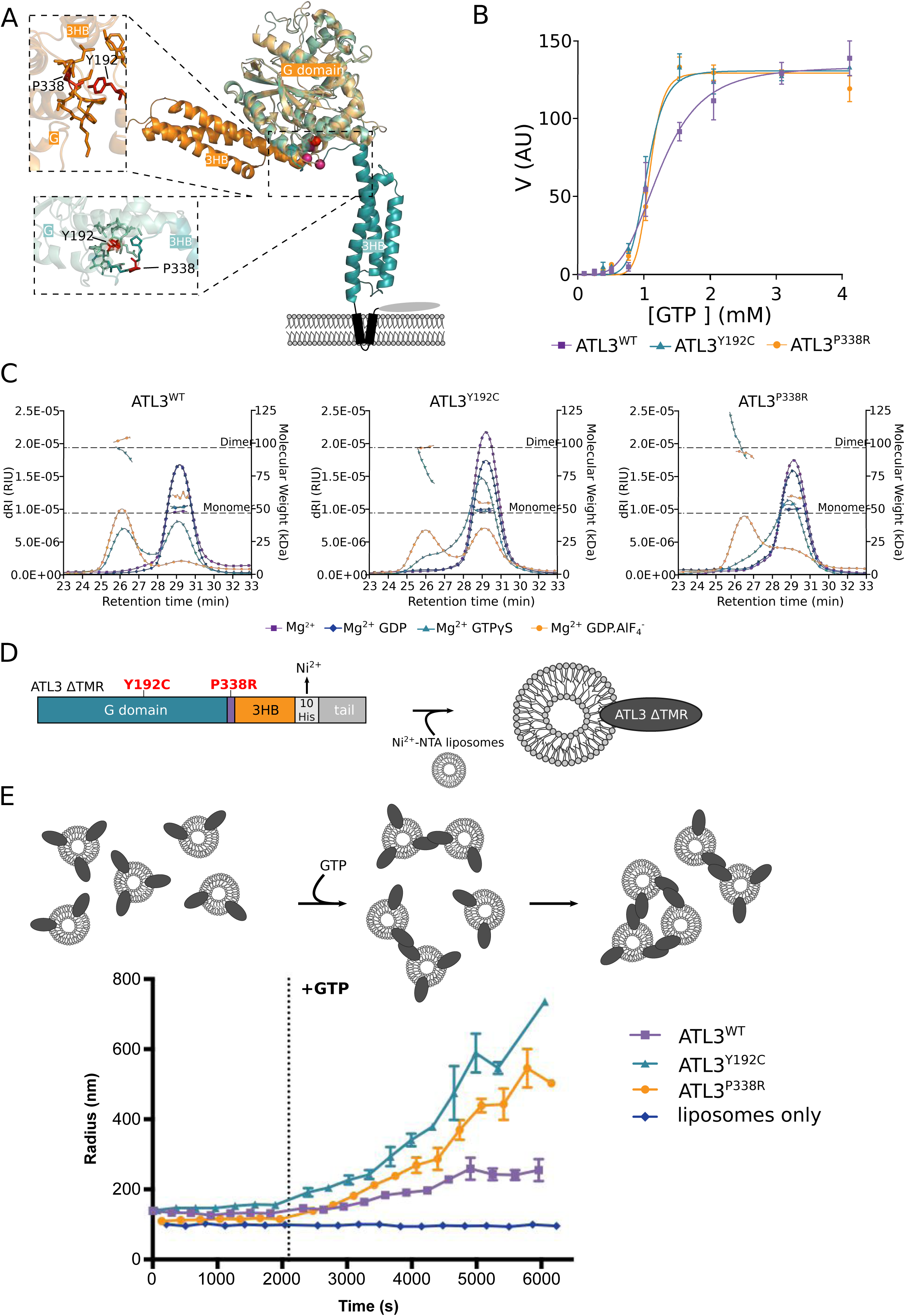
Mutations in ATL3 impair GTP-dependent dimerization and cause excessive membrane tethering *in vitro*. A) The main ATL3 structures are presented in cartoon representation. Residues in the zoom boxes are represented in the “stick” format. The “tight crossover” conformation of ATL3 was based on ATL1 structures (pdb 4IDP) and is shown in light (G-domain) and dark (M-domain) teal. In orange is the recently reported structure of ATL3 in the “extended” conformation (pdb 5VGR). The Cα atoms of residues Y192 and P338 are marked as red spheres in the main structure. The same residues are colored red in the zoom boxes. B) GTPase activity of ATL3 22-442 of WT (squares), Y192C (triangles) and P338R (circles) ATL3 at increasing substrate concentration, fitted to an allosteric sigmoidal curve. C) Oligomerization of purified WT (left), Y192C (middle) and P338R (right) ATL3 22-442 was analyzed using SEC-MALLS. The differential refractive index (dRI, left axis, continuous curve) and molecular weight (right axis, discrete curves above peak maxima) are plotted in function of the SEC retention time. The theoretical mass of the ATL3 monomer and dimer are marked by the dashed horizontal lines. The curves corresponding to the different conditions to which the protein was subjected prior to injection are marked with squares (Mg^2+^), diamonds (Mg^2+^ GDP), triangles (Mg^2+^ GTPγS) and circles (Mg^2+^ GDP.AlF_4_^−^). D) To analyze membrane tethering activity, ATL3 ∆TMR was immobilized on Ni^2+^-NTA containing liposomes through its 10xHis stretch. E) The increase in apparent hydrodynamic radius of the proteoliposomes upon tethering was measured using dynamic light scattering. Tethering was induced by adding 5mM GTP at the indicated time point. See also Fig. S5.

To enable comparative biochemical and biophysical characterization of ATL3^WT^, ATL3^Y192C^, and ATL3^P338R^ we produced and purified recombinant WT and mutant proteins comprising residues 22-442 (Fig. S5A, B). We verified the equivalence of the folding state of the three recombinant proteins by circular dichroism (CD), which revealed mean residual ellipticity (MRE) profiles consistent with the presence of a mix of α-helical and β-strand secondary structural elements in ATLs (Fig. S5C). We first tested whether the HSAN-causing mutations interfere with the GTPase activity of ATL3. We found that the GTPase activity of ATL3^WT^ displayed significant cooperativity, as indicated by the sigmoidal relationship between the enzymatic activity and substrate concentration (Fig. 4B). Such cooperativity is in agreement with dimerization-dependence of the ATL3 GTPase activity. Similar analysis of the catalytic activity of ATL3^Y192C^ and ATL3^P338R^ revealed that these mutant proteins not only retain the competence to hydrolyze GTP, but also maintain the ability to dimerize following GTP binding (Fig. 4B).

Following GTP hydrolysis, the ATL3 dimer is expected to undergo a conformational change whereby both molecules switch from their initial conformation, most likely resembling the extended structure, to a stable crossover dimer (Fig. 4A and Fig. S1) (Bian et al., 2011; Byrnes and Sondermann, 2011; Byrnes et al., 2013; Hu and Rapoport, 2016; Liu et al., 2015; O’Donnell et al., 2017; Saini et al., 2014). *In vitro*, these stable dimers form in the presence of the non-hydrolyzable GTP analog GTPɣS and the transition state analog GDP.AlF_4_^−^ (Morin-Leisk et al., 2011; Saini et al., 2014; Winsor et al., 2017). To examine oligomerization of the ATL3 22-442 proteins in solution, we performed size-exclusion chromatography (SEC) coupled to multi-angle laser light scattering (MALLS). While the SEC retention times correlate with the hydrodynamic volume of the particles, static light scattering allows molecular weight determination regardless of shape (Table S1) and therefore the possible oligomerization state of a given protein. We found that in the absence of nucleotide, all three proteins displayed the same SEC-MALLS profile, consistent with monomeric states averaging 48.1 kDa compared to the theoretical MW of 47.9 kDa (Fig. 4C and Table S1). In the presence of GDP, these profiles remained unchanged (Fig. 4C and Table S1). However, in the presence of GTP analogues, ATL3^WT^ showed a clear tendency to dimerize, corroborating earlier findings (Bian et al., 2011; Byrnes and Sondermann, 2011; Byrnes et al., 2013; O’Donnell et al., 2017). At equilibrium, the presence of GTPɣS separated ATL3^WT^ into two distinct oligomeric states, with about half of the protein adopting a monomeric form and the remaining half a dimeric state (MW 92.7 kDa) (Fig. 4C and Table S1). GDP.AlF_4_^−^ further shifted this equilibrium towards the dimeric species, reducing the monomeric form to about 30% (Fig. 4C and Table S1). Markedly, this oligomerization propensity was severely compromised in the case of ATL3^Y192C^ and ATL3^P338R^. In the presence of GTPɣS, only a limited amount of ATL3^Y192C^ and ATL3^P338R^ was found in the dimer fraction (~20 %) (Fig. 4C and Table S1). While GDP.AlF_4_^−^ again shifted the equilibrium towards dimers, it did so in decreased proportions with only 40% and 50% dimers for ATL3^Y192C^ and ATL3^P338R^, respectively (Fig. 4C and Table S1). Thus, we conclude that the mutations in ATL3 imprint a drastic reduction in the capacity of ATL3 to adopt a stable dimeric state following GTP hydrolysis. Since stable crossover dimer formation is a prerequisite for ATL-mediated membrane fusion (Saini et al., 2014; Winsor et al., 2017), this also explains our earlier observations on defects in ER tubule fusion in cells expressing the pathogenic variants in ATL3 (Fig. 3A,B).

### Mutations in ATL3 result in excessive ER tubule tethering

While being essential for membrane fusion, several studies have shown that stable crossover dimer formation is dispensable for membrane tethering (Saini et al., 2014; Winsor et al., 2017). Therefore, to test for the membrane tethering capacity of ATL3^Y192C^ and ATL3^P338R^, we replaced the transmembrane region of ATL3 with 10 histidines (termed ATL3

∆TMR), which allows binding of the protein to Ni^2+^-NTA containing liposomes (Fig. 4D and Fig. S5D). Since the absence of transmembrane helices of ATL prevents membrane fusion (Liu et al., 2012; Saini et al., 2014), dynamic light scattering (DLS) can be used to analyze the tethering capacity of WT or mutant ATL3 ∆TMR *in vitro* by measuring the apparent increase in size of the proteoliposomes upon addition of GTP (Fig. 4E) (Liu et al., 2015). Indeed, ATL3^WT^ proteoliposomes showed an increase in the mean radius upon the addition of GTP, reaching a plateau around 255nm indicating steady-state tethering between a subset of liposomes at each given time point (Fig. 4E). Unexpectedly, both disease-causing mutations in ATL3 did not only retain liposome-tethering activity, in fact they appeared hyperactive, resulting in a much more pronounced accumulation of tethered liposomes after the addition of GTP (Fig. 4E). These data suggest that while the mutant proteins fail to attain a stable crossover dimer in solution (Fig. 4C), their membrane-tethering capacity is unaffected, showing that they do establish dimers in *trans* when associated with liposomes. Moreover, the defect caused by these mutations results in hyperactive tethering compared to ATL3^WT^, causing an accumulation of tethered liposomes *in vitro* (Fig. 4E).

To address whether this observation was relevant in cells expressing mutant ATL3, we applied focused-ion beam scanning electron microscopy (FIBSEM) which allowed us to resolve the ultrastructure of the ER in large volumes at nanometer-scale resolution (Kremer et al., 2015). Volume-EM images of cells overexpressing ATL3^WT^ revealed that the ER in these cells exhibits a typical morphology hallmarked by an elaborate network of sheets (Fig. 5 upper panel and Video 4). Analysis of a similar region in the periphery of a HeLa cell expressing ATL3^Y192C^ confirmed our earlier findings that the ER is sparsely connected in these cells and consists of long unbranched tubules (Fig. 5 middle panel and Video 5). In addition, we observed several tubules that were attached laterally without the presence of a junction, as indicated by the differently colored ER tubules in our 3D reconstruction (Fig. 5 middle panel and Video 5). This became even clearer when analyzing the perinuclear region in the same cell expressing ATL3^Y192C^. The collapsed ER -as shown in Fig. 1- consisted of a severely anomalous tangle of mostly ER tubules interspersed with a few ER sheets (Video 6). Reconstruction of a part of the ER tangle in this perinuclear region clearly showed the presence of long unbranched tubules that appear tethered laterally without being interconnected, again underscoring that mutant ATL3 is still capable of mediating membrane tethering, but not ER tubule fusion (Fig. 5 lower panel and Video 7). In conclusion, in agreement with the *in vitro* data, the loss of fusogenic capacity in these proteins is accompanied by excessive membrane tethering that has a major impact on ER morphogenesis in cells expressing mutant ATL3.

**Fig. 5:**
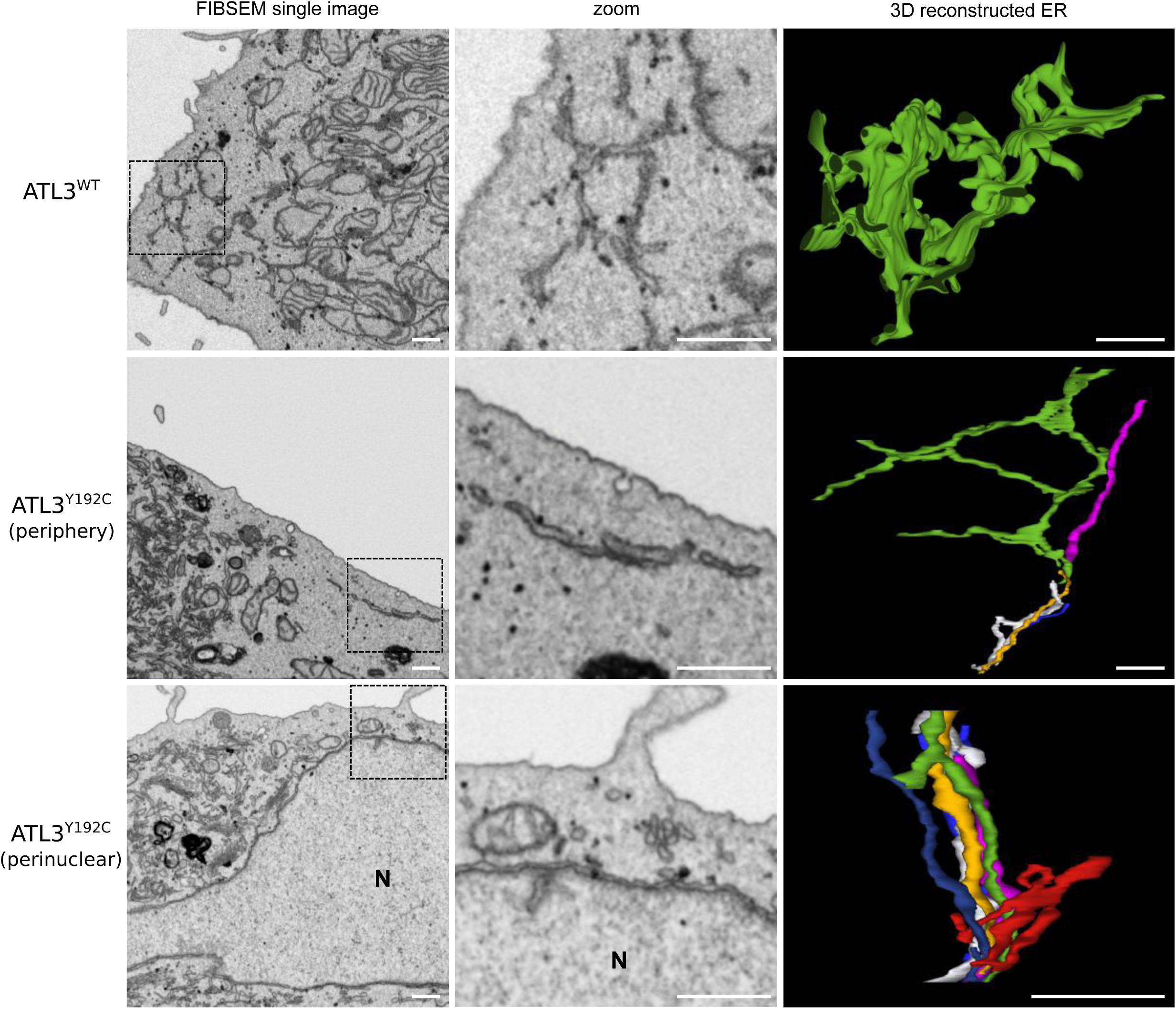
Mutations in ATL3 result in aberrant tethering of unbranched tubules. High-resolution volume electron microscopy of HeLa cells expressing WT (upper row) or mutant (middle and lower rows) ATL3 was performed using focused ion-beam scanning electron microscopy (FIBSEM). A representative section of each cell is shown on the left and a higher magnification of the marked area is shown in the middle column. A 3D model obtained by manual segmentation of a fraction of the ER (as marked in color on the FIBSEM slice) and reconstruction using IMOD is shown on the right. Differently colored tubules indicate that these are discontinuous within the reconstructed volume. See also Videos 4-7. Scale bars = 500nm. N = nucleus.

### Aberrant ER membrane tethering in HSAN patient fibroblasts and neurons

So far, all our *in cellulo* data were generated from cells overexpressing WT or mutant ATL3. To exclude any possible artefact due to overexpression, we derived skin fibroblasts from a patient carrying the ATL3^Y192C^ mutation (Fig. 6). Staining for endogenous KDEL in these cells revealed discernable ER clustering, particularly in the perinuclear region (Fig. 6A). This became even more apparent using transmission electron microscopy. While control fibroblasts displayed a highly connected and well-spread ER, we frequently observed clusters of parallel-oriented unbranched ER membranes in the patient fibroblasts (Fig. 6B and Fig. S6). As expected from the TEM images, FIBSEM analysis of these patient cells indeed revealed several regions of anomalous tethering of ER membranes (Fig. 6C). These observations in HSAN patient-derived cells therefore confirm that aberrant ER tubule tethering caused by mutations in ATL3 is relevant also in conditions of endogenous ATL3 expression.

**Fig. 6:**
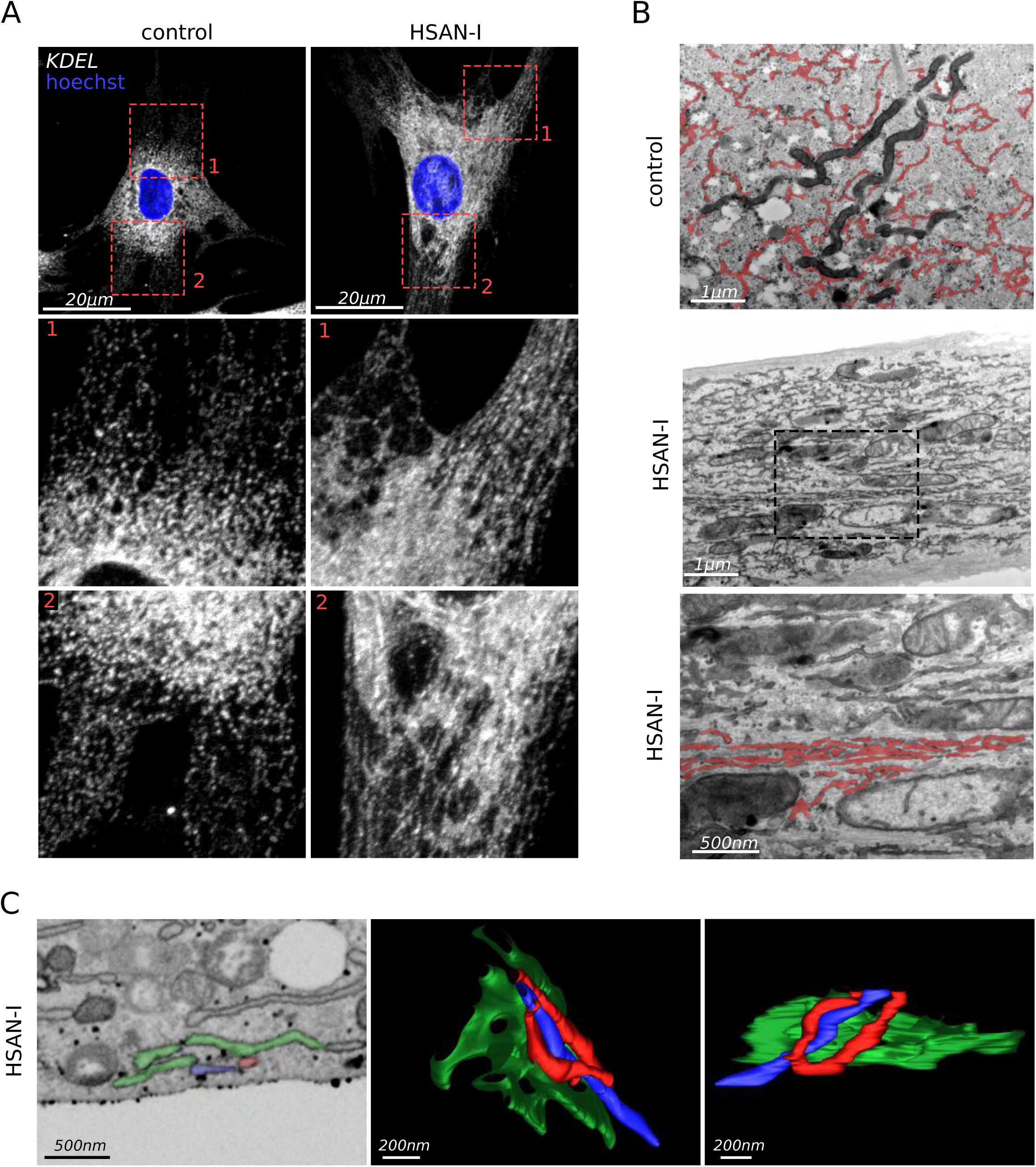
ER tubule clustering is present in HSAN patient fibroblasts. A) Human skin fibroblasts of control or HSAN-I individuals, immunostained for endogenous KDEL as an ER marker. Hoechst nuclear stain is shown in blue. Scale bar = 20µm. Higher magnifications of the boxed areas are shown. B) TEM images of a control fibroblast and a patient fibroblast. The ER in the control cell is marked in red. A higher magnification of the indicated area of the patient cell is shown below, with an example of tangled ER marked in red. The same image without the segementation is shown in Fig. S6. C) FIBSEM imaging of a patient fibroblast. A representative section is shown on the left, and two different angles of a 3D model of the marked ER are shown on the right.

To assess whether this tethering is relevant for neurodegeneration, we expressed WT or mutant ATL3 in primary mouse cortical neurons using lentiviral transduction (Fig. 7). In these cells, the ER marked by ATL3-GFP was found to spread from a dense structure in the soma throughout the neurites regardless of whether WT or mutant ATL3 was expressed, indicating that the mutations do not prevent the generation of neurites or the spread of the ER therein (Fig. 7A). However, in the neurons expressing ATL3^Y192C^ or ATL3^P338R^, we observed a higher ER density within the soma compared to cells expressing ATL3^WT^, similar to what we observed in COS-7 or HeLa cells (Fig. 1 and Fig. 5). While such dense ER clusters (Fig. 7A, arrowheads) were only observed in less than 10% of neurons overexpressing ATL3^WT^, approximately 70% of cells expressing mutant ATL3 exhibited this phenotype, showing that this is a major hallmark of HSAN-causing mutations in ATL3 (Fig. 7B). To show that these structures indeed represent an altered distribution of ATL3-GFP throughout the cells, we measured relative GFP intensities along single neurites starting from the center of the soma, which confirmed that neurons expressing mutant ATL3-GFP retained a larger fraction of the total GFP signal within this perinuclear ER structure compared to ATL3^WT^-expressing cells (Fig. 7C). In order to resolve the ultrastructure of the ER in these clusters, we performed FIBSEM imaging on these neurons using correlative light electron microscopy (CLEM) to identify the neurons expressing GFP-tagged ATL3 (Fig. 7D). Similar to what we observed in patient fibroblasts, also the neurons expressing ATL3^Y192C^ showed distinct regions of tethered ER within the high-density ER structures that were observed by light microscopy (Fig. 7E and Video 8-9). Such clusters were never observed in primary cortical neurons overexpressing ATL3^WT^ (Fig. S7). In summary, these data show that both in patient fibroblasts and in neuronal cells mutations of ATL3 lead to a partial collapse of the ER due to excessive membrane tethering, further underscoring the physiological relevance of our findings.

**Fig. 7:**
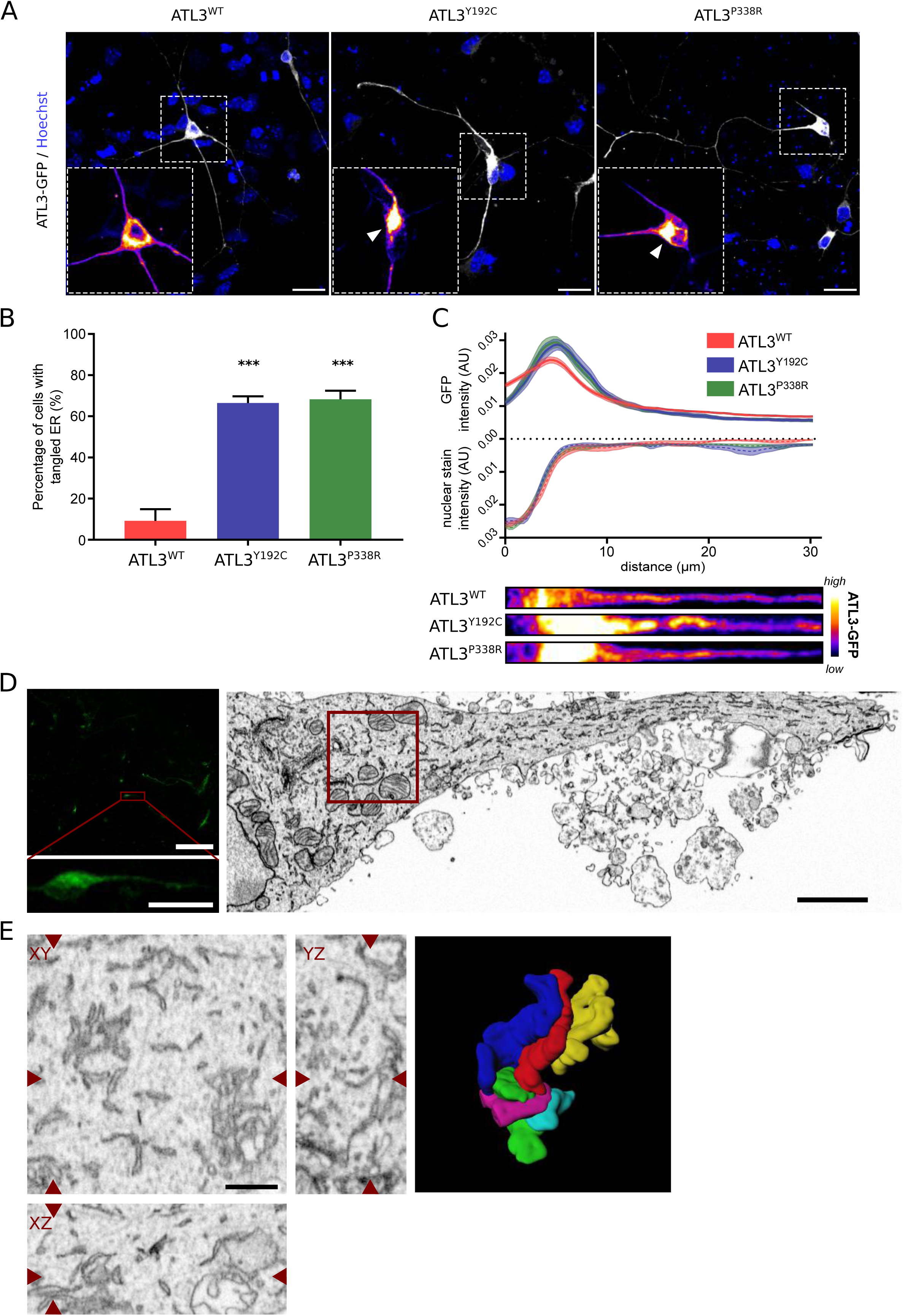
ER tangling is observed in primary cortical neurons expressing mutant ATL3. A) Mouse primary cortical neurons were transduced with ATL3-GFP and Hoechst nuclear stain is shown in blue. Higher magnifications of ATL3-GFP in the boxed areas are shown in a fluorescence intensity color code as indicated in C. Arrow heads mark dense ER clusters in the mutant neurons. Scale bar = 20µm. B) The fraction of cells expressing WT or mutant ATL3 displaying ER tangling (as marked by arrowheads in A) was determined in 3 independent cortical neuron isolations (n=39-53 cells). Mean ± SEM is plotted. ***: P < 0.001 in a one-way Anova test. C) Normalized GFP and nuclear stain fluorescence intensity are plotted in function of the position along a line segment that traces a single neurite and starts in the center of the nucleus. A ‘straightened’ version of such a 30µm neurite tracing from the cells shown in A, is shown for each genotype below the graph. Mean ± SEM is plotted (n=30-35 cells). D) Primary cortical neurons expressing GFP-tagged ATL3^Y192C^ (left). Scale bars = 100 µm in the overview image and 20µm in the zoomed region. The same neuron was identified in the electron microscope (right, scale bar = 2µm). E) A cluster of tangled ER membranes was identified in the perinuclear area marked in D. Also XZ and YZ projections of the positions marked by arrowheads are shown. Tangled ER membranes were segmented and reconstructed in 3D. Scale bar = 500nm. Volume-EM for a cortical neuron expressing ATL3^WT^ is shown in Fig. S7.

## DISCUSSION

In this study, we have provided new insights into how disease-related mutations in ATL3 interfere with the protein’s functional cycle required for membrane fusion. Importantly, we show that such functional defects impact the structure and integrity of the ER both in cells overexpressing pathogenic variants of ATL3 and in patient-derived fibroblasts. Indeed, the formation of new junctions through ER tubule fusion mediated by the Atlastin GTPases is essential for the continuous remodeling of the ER, defects in which cause axonal degeneration in diseases such as HSAN and HSP (Fischer et al., 2014; Guelly et al., 2011; Kornak et al., 2014; Zhao et al., 2001). Here, we show that in addition to a loss of fusion activity, the HSAN-causing ATL3 variants cause lateral tethering and tangling of ER membranes.

Recent structural and biochemical studies on human ATL1-3 and *Drosophila* ATL (dATL) have proposed a model for ATL-mediated membrane fusion in which the G domains of ATL1 dimerize *in trans* upon GTP binding (Fig. S1A), and that GTP hydrolysis promotes the formation of a tight crossover dimer (Fig. S1B), bringing both membranes in close apposition and thereby driving membrane fusion (Bian et al., 2011; Byrnes and Sondermann, 2011; Byrnes et al., 2013; Liu et al., 2015; O’Donnell et al., 2017; Saini et al., 2014; Winsor et al., 2017). Our data now show that the disease-causing ATL3 variants do not interfere with the initial steps of the functional cycle leading up to G domain dimerization, nor do they affect the allosteric properties (Fig. 4). However, the mutant proteins do show a different behavior in the steps following initial dimerization. The ATL3^Y192C^ and ATL3^P338R^ mutant proteins suffer from a reduced dimerization propensity in the presence of GTPɣS or GDP+AlF_4_^−^ (Fig. 4), suggesting that the mutants fail to establish the stable crossover dimer conformation following GTP hydrolysis. As this stable dimer is an absolute prerequisite to cross the energy barrier required for merging lipid bilayers (Winsor et al., 2017), failure to attain this conformation offers an attractive explanation for the inability of the mutant ATL3 to support membrane fusion, as reflected by the ER network connectivity deficit observed in COS-7 cells (Fig. 1-3).

In contrast to fusion, the preceding membrane tethering step does not depend on the stable crossover dimer (Liu et al., 2015; Morin-Leisk et al., 2011; Saini et al., 2014; Winsor et al., 2017). Interestingly, one of the most striking ER defects we observed in cells expressing HSAN-causing mutations in ATL3, was the presence of extensive ER membrane tethering as revealed by volume-EM. This unique phenotype was characterized by the presence of bundles of ER tubules running in parallel over a large number of FIBSEM slices (Fig. 5 and Videos 6, 7). This aberrant tethering occurred mainly in a collapsed, high-density region of the ER near the nucleus in cell lines. Although we currently do not have direct structural data to rationalize the underlying mechanism, we hypothesize that HSAN-causing mutations in ATL3, located in the interface between the G and the 3HB domain, prevent the protein from proceeding through its functional cycle following *trans*-dimerization and GTP hydrolysis. Rather than completing their cycle followed by dissociation of the dimer, the proteins seem stalled or blocked in the membrane tethering stage, causing excessive membrane tethering in cell lines, fibroblasts and neurons (Fig. 5-7). These interactions were not retrieved in the SEC-MALLS experiments, indicating that they are transient in solution and might require further stabilization by membrane association and the 2-dimensional spatial confinement of the membrane (Winsor et al., 2017). Indeed, the *in vitro* tethering assay confirmed that both disease-causing mutations in ATL3 promote excessive liposome tethering compared to the WT protein (Fig. 4).

Together, our findings are in agreement with a model in which GTP binding and hydrolysis drives ER membrane tethering through ATL trans-dimerization. Subsequent membrane fusion relies on the formation of a stable crossover dimer bringing the membranes in close apposition, resulting in a new three-way junction. While the HSAN-causing mutations in ATL3 do not abrogate GTP binding or hydrolysis, the mutant proteins fail to completely execute the subsequent conformational changes required to form a stable crossover dimer, rendering the mutant ATL3 fusion-deficient. Underscoring the relevance of this stable crossover conformation to reset the protein for a new round of tethering and fusion (O’Donnell et al., 2017), the inability of both mutants to proceed to this step of the ATL functional cycle causes aberrant membrane tethering that leads to a partial collapse of the ER network, strongly affecting ER dynamics.

ATL1 is the predominant isoform in the nervous system (Kornak et al., 2014; Rismanchi et al., 2008), and also the Atlastin with the highest intrinsic fusogenic capacity (Hu et al., 2015; O’Donnell et al., 2017). Therefore, a defect in ER fusion is thought to underlie the HSP phenotype in most of the patients carrying mutations in ATL1 (Ulengin et al., 2015). ATL3 on the other hand was found to be a much weaker fusogen (O’Donnell et al., 2017), and loss of ATL3 has only marginal effects on overall ER morphology (Hu et al., 2015). Therefore, in the case of the dominant missense mutations in ATL3 causing HSAN, loss of ER fusion is far less likely to be the main factor contributing to disease. In contrast, a partial collapse of the ER network caused by the excessive membrane-tethering in the ATL3^Y192C^ and ATL3^P338R^ mutants might have a much higher impact on neuronal functioning. In agreement with this, ER tangling was also detected in patient fibroblast where the mutant ATL3 is expressed at physiologically relevant levels (Fig. 6). Similarly, ER tangling was observed in neurons expressing mutant ATL3, affecting the distribution of the ER within these cells (Fig. 7). We hypothesize that ER tethering disturbs normal ER membrane dynamics, and most likely interferes with the organization of different ER subdomains playing various roles in neuronal homeostasis (English and Voeltz, 2013). In agreement with our hypothesis, no measurable defect in ER fusion was found for several HSP-causing variants in ATL1, including the most commonly identified R239C mutation (Ulengin et al, 2015), indicating that defects other than loss of fusion contribute to neurodegeneration in these patients. Since little is currently known about the function of the ER within axons, studying the effect of disease-causing mutations in ATLs and other ER-shaping proteins (Hübner & Kurth, 2014) may shed more light on the role of axonal ER in neuronal homeostasis and disease.

## EXPERIMENTAL PROCEDURES

For full experimental details, please see supplemental information.

### Plasmids, cell lines and transfections

All ATL3-expressing vectors were created using Gateway recombination system (ThermoFisher Scientific). The pCI-mRubyKDEL vector was a gift from Dr. Jörg Wiedenmann (Kredel et al., 2009). The PA-GFP-KDEL vector was a gift from Dr. Vicky Claire Jones (Jones et al., 2009).

Transient transfections of HeLa and COS-7 were performed using Lipofectamine-LTX (ThermoFisher Scientific). Mouse cortical neurons were extracted from E15.5 C57BL/6 embryos and lentiviral transduction was used to express ATL3-GFP. Primary dermal fibroblasts were generated from a patient with a c.575A>G (p.Tyr192Cys) mutation in the ATL3 gene.

### Immunofluorescence microscopy

HeLa, COS-7, mouse cortical neurons and human skin fibroblasts were cultured, PFA-fixed and immunostained according to standard protocols. Imaging was performed on a Zeiss LSM700 confocal microscope. Image analysis was done in ImageJ or CellProfiler software (Kamentsky et al., 2011; Schindelin et al., 2012).

### ER morphology and fusion success rate

COS-7 cells transfected with ATL3-GFP and mRuby-KDEL were imaged using spinning disk confocal microscopy at 1 frame per sec. ER sparsity and junction density were measured on the first frame using ImageJ/FIJI (Schindelin et al., 2012; Schneider et al., 2012). Tubule fusion efficiency was scored as the proportion of successful fusion events over total extending tubule tip contact events.

### Photo-activatable GFP imaging

COS-7 cells transfected with ATL3-mCherry and PA-GFP-KDEL were imaged on a Zeiss LSM700 microscope. PA-GFP-KDEL was activated in a perinuclear ER region using the 405nm laser at 100% and imaged for 90s using the 488nm laser at 1 frame per second. Fluorescence intensities were measured using ImageJ (Schindelin et al., 2012; Schneider et al., 2012) and data analysis and curve fitting was performed in R (http://www.R-project.org/).

### Transmission and Volume electron microscopy

Hela cells and primary cortical neurons were seeded in 35mm imaging dishes with a gridded coverslip (MatTek). The position of the cells of interest in the alphanumerically labelled grid was determined by imaging GFP in living cells. After fixation cells were further processed for imaging on the Zeiss Auriga Crossbeam system as described (Knott et al., 2011). FIBSEM imaging at a 5×5×5 nm voxel size was performed as previously described (Kremer et al., 2015). Image stacks were registered and processed with ImageJ and ER reconstructions were made using 3D-MOD software.

Transmission electron microscopy on human skin fibroblasts was performed according to routine procedures, including embedding in Spurr’s resin, ultrathin sectioning (Leica EM UC7) and imaging using a JEM 1400plus transmission electron microscope (JEOL).

### GTPase activity

Recombinant WT and mutant ATL3 were expressed in E. coli with a thrombin-cleavable HIS(6)-tag using standard approaches. GTPase activity was measured using the EnzCheck phosphate assay kit (ThermoFisher Scientific).

### SEC-MALLS

Size exclusion chromatography - Multi-Angle Laserlight Scattering (SEC-MALLS) experiments were performed on a sequential setup encompassing Degasys degasser (Gynkotek), 1260 Infinity HPLC pump (Agilent), Superdex 200 Increase 10/300 GL SEC column (GE Healthcare), SPD-10A UV-VIS detector (Shimadzu), miniDAWN TREOS light scattering detector (Wyatt) and Optilab T-rEX refraction index detector (Wyatt). Prior to injection, protein samples (100 µL at 1 mg/ml) were incubated for 90’ at room temperature with the indicated nucleotides. Data were recorded and analyzed using the ASTRA software package (Wyatt, v6.1).

### Liposome tethering

Timecourse dynamic light scattering measurements were performed at 37°C in a DynaPro plate reader-II (Wyatt technology) using reactions consisting of 120 µM Ni^2+^-NTA containing liposomes and 1 µM ATL3 ∆TMR. After 30 minutes, 5mM GTP was added to induce liposome tethering.

## Author contributions

Conceptualization, M.K., B.A., L.A.-S. and S.J.; Methodology, M.K., S.D., B.A., L.A.-S., S.N.S. and S.J.; Investigation, M.K., S.D., B.A., L.A.-S., A.K., S.L., R.D.R. and V.D.W.; Writing-Original draft, M.K., S.D., S.N.S. and S.J.; Writing-Review and editing, M.K., S.D., B.A., L.A.-S., S.N.S., V.T. and S.J.; Funding Acquisition, M.K., V.T., S.J.; Resources, I.K., F-J.M., H.T.M; Supervision, S.L., H.T.M., S.N.S., V.T. and S.J.

## Acknowledgements

M.K. is supported by a predoctoral fellowship of the Fund for Scientific Research (FWO-Flanders, Belgium) and a short term EMBO fellowship. S.D. is supported by a predoctoral fellowship from the Flemish Agency for Innovation and Entrepreneurship (VLAIO). L.A.-S. is an EMBO Long Term fellow and is supported by Marie Curie Actions. F.J.M. is supported by grants from the BMBF (13GW0128A and 01GM1513D) and from the Deutsche Forschungsgemeinschaft (DFG MU 3231/3-1). H.T.M. is funded by the Medical Research Council UK (grant number U105178795). The research of V.T. is supported by the FWO, University of Antwerp, the Hercules Foundation, the “Association Belge contre les Maladies Neuro-Musculaires” (ABMM) and the EU FP7/2007–2013 under grant agreement number 2012-305121 (NEUROMICS). The research program of S.J. is supported by the FWO, Ghent University and GROUP-ID. S.N.S. is supported by the FWO, the Hercules Foundation, and the VIB. The Zeiss Auriga was acquired through a CLEM grant from Minister Ingrid Lieten to the VIB Bio Imaging Core. Special thanks to Elias Adriaenssens, Sonia Bartunkova, Michiel De Bruyne, Isabel Pintelon and Peter Verstraelen for advice and technical support. Also thanks to Chris Guérin for advice in image processing. Thanks to Vicky Claire Jones and Joerg Wiedenmann for providing plasmids. Molecular graphics and analyses were performed with the UCSF Chimera package. Chimera is developed by the Resource for Biocomputing, Visualization, and Informatics at the University of California, San Francisco (supported by NIGMS P41-GM103311).

## VIDEO LEGENDS

Video 1: **Photoactivatable GFP-KDEL**. PA-GFP-KDEL was transiently expressed in COS-7 cells, along with ATL3-mCherry as indicated in Fig. 2. Imaging was performed at 1 frame per second, and 5 frames were recorded prior to photoactivation of the PA-GFP-KDEL in a small region near the nucleus.

Video 2: **Successful tubule fusion.** mRuby-KDEL was transiently transfected in COS-7 cells to label the ER, along with ATL3-GFP as indicated in Fig. 3. To monitor tubule encounter events, live-cell imaging was performed at 1 frame per second. In this example, a new threeway junction is formed upon encounter between the extending tubule at the far left and the opposing tubule.

Video 3: **Failed tubule fusion.** As in Fig. 3 and Video 2, COS-7 cells were transfected with mRuby-KDEL and imaged at 1 frame per second. In this example, the tubule encounter occurring at the center of the view fails to result in a new junction, and the extending tubule eventually retracts.

Video 4: **ER ultrastructure in an ATL3^WT^-expressing HeLa visualized using volume electron microscopy.** Focused-ion beam scanning electron microscopy (FIBSEM) was performed on a HeLa cell expressing WT ATL3-GFP. This video shows a digitally resliced version of the original image stack, and a 3D model of a portion of the ER in green, as in Fig. 5.

Video 5: **ER tubule tangling and loss of network complexity in a HeLa cell expressing ATL3^Y192C^.** Focused-ion beam scanning electron microscopy (FIBSEM) was performed on a HeLa cell expressing Y192C ATL3-GFP. A 3D reconstruction of a portion of the ER is shown as in Fig. 5.

Video 6: **ER tubule tangling and loss of network complexity in a HeLa cell expressing ATL3^Y192C^.** Focused-ion beam scanning electron microscopy (FIBSEM) was performed on a HeLa cell expressing Y192C ATL3-GFP. This video shows a perinuclear region of this cell (Fig. 5), revealing a tangle of ER membranes that represents the collapsed ER seen in Fig. 1.

Video 7: **Extensive ER tubule tangling in a perinuclear region of a HeLa cell expressing ATL3^Y192C^.** Focused-ion beam scanning electron microscopy (FIBSEM) was performed on a HeLa cell expressing Y192C ATL3-GFP. This video shows a perinuclear region of this cell (Fig. 5 and Video 6). 3D reconstruction of the ER reveals extensive lateral tubule tethering (Fig. 5). Different colors indicate tubules that are not continuous within the reconstructed volume.

Video 8: **ER tubule tangling in a primary cortical neuron expressing ATL3^Y192C^.** Focused-ion beam scanning electron microscopy (FIBSEM) was performed on a cortical neuron expressing Y192C ATL3-GFP. This video shows an ER tubule cluster within the soma (Fig. 8).

Video 9: **3D reconstruction of an ER tubule cluster in a primary cortical neuron expressing ATL3^Y192C^.** The tangled ER tubules in Video 8 were segmented for 3D reconstruction. Different colors indicate tubules that are not continuous within the reconstructed volume.

## References

Bian, X., R.W. Klemm, T.Y. Liu, M. Zhang, S. Sun, X. Sui, X. Liu, T.A. Rapoport, and J. Hu. 2011. Structures of the atlastin GTPase provide insight into homotypic fusion of endoplasmic reticulum membranes. Proceedings of the National Academy of Sciences of the United States of America. 108:3976–3981.

Byrnes, L., and H. Sondermann. 2011. Structural basis for the nucleotide-dependent dimerization of the large G protein atlastin-1/SPG3A. PNAS.

Byrnes, L.J., A. Singh, K. Szeto, N.M. Benvin, J.P. O’Donnell, W.R. Zipfel, and H. Sondermann. 2013. Structural basis for conformational switching and GTP loading of the large G protein atlastin. The EMBO journal. 32:369–384.

de Bot, S.T., J.H. Veldink, S. Vermeer, A.R. Mensenkamp, F. Brugman, H. Scheffer, L.H. van den Berg, H.P. Kremer, E.J. Kamsteeg, and B.P. van de Warrenburg. 2013. ATL1 and REEP1 mutations in hereditary and sporadic upper motor neuron syndromes. J Neurol. 260:869–875.

English, A.R., and G.K. Voeltz. 2013. Endoplasmic reticulum structure and interconnections with other organelles. Cold Spring Harb Perspect Biol. 5:a013227.

Fischer, D., M. Schabhuttl, T. Wieland, R. Windhager, T.M. Strom, and M. Auer-Grumbach. 2014. A novel missense mutation confirms ATL3 as a gene for hereditary sensory neuropathy type 1. Brain. 137.

Greenfield, N.J. 2006. Using circular dichroism spectra to estimate protein secondary structure. Nature protocols. 1:2876–2890.

Guelly, C., P.-P.P. Zhu, L. Leonardis, L. Papić, J. Zidar, M. Schabhüttl, H. Strohmaier, J. Weis, T.M. Strom, J. Baets, J. Willems, P. De Jonghe, M.M. Reilly, E. Fröhlich, M. Hatz, S. Trajanoski, T.R. Pieber, A.R. Janecke, C. Blackstone, and M. Auer-Grumbach. 2011. Targeted high-throughput sequencing identifies mutations in atlastin-1 as a cause of hereditary sensory neuropathy type I. American journal of human genetics. 88:99–105.

Hu, J., and T.A. Rapoport. 2016. Fusion of the endoplasmic reticulum by membrane-bound GTPases. Seminars in Cell & Developmental Biology.

Hu, J., Y. Shibata, P.-P.P. Zhu, C. Voss, N. Rismanchi, W.A. Prinz, T.A. Rapoport, and C. Blackstone. 2009. A class of dynamin-like GTPases involved in the generation of the tubular ER network. Cell. 138:549–561.

Hu, X., F. Wu, S. Sun, W. Yu, and J. Hu. 2015. Human atlastin GTPases mediate differentiated fusion of endoplasmic reticulum membranes. Protein & cell. 6:307–311.

Hübner, C.A., and I. Kurth. 2014. Membrane-shaping disorders: a common pathway in axon degeneration. Brain: a journal of neurology. 137:3109–3121.

Jones, V.C., J.J.J. Rodríguez, A. Verkhratsky, and O.T. Jones. 2009. A lentivirally delivered photoactivatable GFP to assess continuity in the endoplasmic reticulum of neurones and glia. Pflugers Archiv: European journal of physiology. 458:809–818.

Kamentsky, L., T.R. Jones, A. Fraser, M.A. Bray, D.J. Logan, K.L. Madden, V. Ljosa, C. Rueden, K.W. Eliceiri, and A.E. Carpenter. 2011. Improved structure, function and compatibility for CellProfiler: modular high-throughput image analysis software. Bioinformatics. 27:1179–1180.

Knott, G., S. Rosset, and M. Cantoni. 2011. Focussed ion beam milling and scanning electron microscopy of brain tissue. J Vis Exp:e2588.

Kornak, U., I. Mademan, M. Schinke, M. Voigt, P. Krawitz, J. Hecht, F. Barvencik, T. Schinke, S. Gießelmann, F.T. Beil, A. Pou-Serradell, J.J. Vílchez, C. Beetz, T. Deconinck, V. Timmerman, C. Kaether, P. De Jonghe, C.A. Hübner, A. Gal, M. Amling, S. Mundlos, J. Baets, and I. Kurth. 2014. Sensory neuropathy with bone destruction due to a mutation in the membrane-shaping atlastin GTPase 3. Brain: a journal of neurology. 137:683–692.

Kredel, S., F. Oswald, K. Nienhaus, K. Deuschle, C. Röcker, M. Wolff, R. Heilker, G.U. Nienhaus, and J. Wiedenmann. 2009. mRuby, a bright monomeric red fluorescent protein for labeling of subcellular structures. PloS one. 4.

Kremer, A., S. Lippens, S. Bartunkova, B. Asselbergh, C. Blanpain, M. Fendrych, A. Goossens, M. Holt, S. Janssens, M. Krols, J.C. Larsimont, C. Mc Guire, M.K. Nowack, X. Saelens, A. Schertel, B. Schepens, M. Slezak, V. Timmerman, C. Theunis, V.A.N.B. R, Y. Visser, and C.J. Guerin. 2015. Developing 3D SEM in a broad biological context. Journal of microscopy. 259:80–96.

Larkin, M.A., G. Blackshields, N.P. Brown, R. Chenna, P.A. McGettigan, H. McWilliam, F. Valentin, I.M. Wallace, A. Wilm, R. Lopez, J.D. Thompson, T.J. Gibson, and D.G. Higgins. 2007. Clustal W and Clustal X version 2.0. Bioinformatics (Oxford, England). 23:2947–2948.

Liu, T.Y., X. Bian, F.B. Romano, T. Shemesh, T.A. Rapoport, and J. Hu. 2015. Cis and trans interactions between atlastin molecules during membrane fusion. Proceedings of the National Academy of Sciences of the United States of America. 112:60.

Liu, T.Y., X. Bian, S. Sun, X. Hu, R.W. Klemm, W.A. Prinz, T.A. Rapoport, and J. Hu. 2012. Lipid interaction of the C terminus and association of the transmembrane segments facilitate atlastin-mediated homotypic endoplasmic reticulum fusion. Proceedings of the National Academy of Sciences of the United States of America. 109:54.

McCorquodale, D.S., 3rd, U. Ozomaro, J. Huang, G. Montenegro, A. Kushman, L. Citrigno, J. Price, F. Speziani, M.A. Pericak-Vance, and S. Zuchner. 2011. Mutation screening of spastin, atlastin, and REEP1 in hereditary spastic paraplegia. Clin Genet. 79:523–530.

Morin-Leisk, J., S.G. Saini, X. Meng, A.M. Makhov, P. Zhang, and T.H. Lee. 2011. An intramolecular salt bridge drives the soluble domain of GTP-bound atlastin into the postfusion conformation. J Cell Biol. 195:605–615.

Moss, T.J., C. Andreazza, A. Verma, A. Daga, and J.A. McNew. 2011. Membrane fusion by the GTPase atlastin requires a conserved C-terminal cytoplasmic tail and dimerization through the middle domain. Proceedings of the National Academy of Sciences of the United States of America. 108:11133–11138.

Namekawa, M., M.P.P. Muriel, A. Janer, M. Latouche, A. Dauphin, T. Debeir, E. Martin, C. Duyckaerts, A. Prigent, C. Depienne, A. Sittler, A. Brice, and M. Ruberg. 2007. Mutations in the SPG3A gene encoding the GTPase atlastin interfere with vesicle trafficking in the ER/Golgi interface and Golgi morphogenesis. Molecular and cellular neurosciences. 35:1–13.

O’Donnell, J.P., R.B. Cooley, C.M. Kelly, K. Miller, O.S. Andersen, R. Rusinova, and H. Sondermann. 2017. Timing and Reset Mechanism of GTP Hydrolysis-Driven Conformational Changes of Atlastin. Structure. 25:997–1010 e1014.

Orso, G., D. Pendin, S. Liu, J. Tosetto, T.J. Moss, J.E. Faust, M. Micaroni, A. Egorova, A. Martinuzzi, and J.A. McNew. 2009. Homotypic fusion of ER membranes requires the dynamin-like GTPase atlastin. Nature. 460:978–983.

Powers, R.E., S. Wang, T.Y. Liu and T.A. Rapoport. 2017. Reconstitution of the tubular endoplasmic reticulum network with purified components. Nature. 543:257–260.

Rismanchi, N., C. Soderblom, J. Stadler, P.P. Zhu, and C. Blackstone. 2008. Atlastin GTPases are required for Golgi apparatus and ER morphogenesis. Hum Mol Genet. 17:1591–1604.

Saini, S.G., C. Liu, P. Zhang, and T.H. Lee. 2014. Membrane tethering by the atlastin GTPase depends on GTP hydrolysis but not on forming the cross-over configuration. Molecular biology of the cell. 25:3942–3953.

Schindelin, J., I. Arganda-Carreras, E. Frise, V. Kaynig, M. Longair, T. Pietzsch, S. Preibisch, C. Rueden, S. Saalfeld, B. Schmid, J.Y. Tinevez, D.J. White, V. Hartenstein, K. Eliceiri, P. Tomancak, and A. Cardona. 2012. Fiji: an open-source platform for biological-image analysis. Nat Methods. 9:676–682.

Schneider, C.A., W.S. Rasband, and K.W. Eliceiri. 2012. NIH Image to ImageJ: 25 years of image analysis. Nat Methods. 9:671–675.

Ulengin, I., J.J. Park, and T.H. Lee. 2015. ER network formation and membrane fusion by atlastin1/SPG3A disease variants. Mol Biol Cell. 26:1616–1628.

Webb, B., and A. Sali. 2016. Comparative Protein Structure Modeling Using MODELLER. In Current Protocols in Protein Science. Vol. 86. John Wiley & Sons, Inc., Hoboken, NJ, USA. 2.9.1–2.9.37.

Westrate, L.M., J.E. Lee, W.A. Prinz, and G.K. Voeltz. 2015. Form follows function: the importance of endoplasmic reticulum shape. Annual review of biochemistry. 84:791–811.

Winsor, J., D.D. Hackney, and T.H. Lee. 2017. The crossover conformational shift of the GTPase atlastin provides the energy driving ER fusion. J Cell Biol.

Zhao, X., D. Alvarado, S. Rainier, R. Lemons, P. Hedera, C.H. Weber, T. Tukel, M. Apak, T. Heiman-Patterson, L. Ming, M. Bui, and J.K. Fink. 2001. Mutations in a newly identified GTPase gene cause autosomal dominant hereditary spastic paraplegia. Nat Genet. 29:326–331.

